# Structural and mechanistic insights into inhibition of the type I-F CRISPR-Cas surveillance complex by AcrIF4

**DOI:** 10.1101/2022.08.30.504949

**Authors:** Zhengyu Gao, Laixing Zhang, Zihao Ge, Hao Wang, Yourun Yue, Zhuobing Jiang, Xin Wang, Chenying Xu, Yi Zhang, Maojun Yang, Yue Feng

## Abstract

CRISPR-Cas system provides prokaryotes with protection against mobile genetic elements (MGEs) such as phages. In turn, phages deploy anti-CRISPR (Acr) proteins to evade this immunity. AcrIF4, an Acr targeting the type I-F CRISPR-Cas system, has been reported to bind the crRNA-guided surveillance (Csy) complex. However, it remains controversial whether AcrIF4 inhibits target DNA binding to the Csy complex. Here, we present structural and mechanistic studies into AcrIF4, exploring its unique anti-CRISPR mechanism. While the Csy-AcrIF4 complex displays decreased affinity for target DNA, it is still able to bind the DNA. Structural and functional analyses of the Csy-AcrIF4-dsDNA complex revealed that AcrIF4 binding prevents rotation of the helical bundle (HB) of Cas8f induced by dsDNA binding, therefore resulting in failure of Cas2/3 recruitment and DNA cleavage. Overall, our study provides an interesting example of attack on the nuclease recruitment event by an Acr, but not conventional mechanisms of blocking binding of target DNA.

## Introduction

Clustered regularly interspaced short palindromic repeats (CRISPRs) and CRISPR-associated (Cas) genes in prokaryotes encode an adaptive immune system that provides protection against the invasive mobile genetic elements (MGEs) such as phages (1,2). The CRISPR-Cas system is classified into two broad classes and further divided into six types (I–VI), based on a wide variation in the protein composition among different types (3). Class 1 systems (types I, III, and IV) account for ∼90% of the CRISPR-Cas systems observed in nature, which depend on CRISPR RNA (crRNA)-guided multi-subunit complexes to recognize foreign nucleic acids. Class 2 systems (types II, V and VI) deploy a single multi-domain protein that serves as a crRNA-guided effector nuclease to target foreign DNA/RNA. Although CRISPR-Cas systems are diverse in their protein composition, they share similar working stages. In their immunity mechanism, integrating short foreign DNA fragments into the host CRISPR locus is used for recognizing and destroying previously encountered non-self nucleic acid sequences, which is the most distinctive feature of CRISPR-Cas system as an adaptive immune system (4).

To evade this bacterial immunity, phages have evolved protein inhibitors called anti-CRISPR (Acr) proteins (5). These proteins differ greatly in their sequence and structure, providing diverse inhibitory mechanisms that act on different components of the CRISPR-Cas system and at distinct stages of the CRISPR-Cas immunity (6). The type I-F CRISPR-Cas system encodes a ∼350 kDa crRNA-guided ribonucleoprotein complex (named the Csy complex), which comprises four kinds of protein (one Cas5f, one Cas8f, one Cas6f and six Cas7f subunits). After the Csy complex recognizes target DNA, the trans-acting nuclease-helicase Cas2/3 will be recruited and degrades the DNA. Up to now, 24 Acrs have been identified to target the type I-F CRISPR-Cas system, out of which both structures and inhibition mechanisms of 14 Acr proteins have been determined, including AcrIF1/2/3/5/6/7/8/9/10/11/13/14/23/24. Among these, all Acrs except AcrIF3/5/23 act by inhibiting target dsDNA binding (7-9), mainly in two approaches, either inhibiting hybridization between target DNA and crRNA or imitating DNA substrates. There are also several novel mechanisms based on this canonical one. AcrIF11 is an ADP-ribosyltransferase that modifies the key Csy residue responsible for target DNA recognition (10). AcrIF9/14/24 are dual functional Acrs with an ability to not only inhibit DNA-crRNA hybridization but also induce non-specific DNA binding to the Csy complex (11-13). AcrIF3 and AcrIF5 do not inhibit target DNA binding by the Csy complex, but prevent subsequent Cas2/3 recruitment. Target dsDNA binding by the Csy complex will induce a ∼180° rotation of the helical bundle (HB) of Cas8f subunit, which exposes an α helix responsible for Cas2/3 recruitment (7). Both AcrIF3 and AcrIF5 exploit this process to achieve their inhibition but function in different ways. AcrIF5 specifically targets the dsDNA-bound Csy complex to reposition the Cas8f HB (9), while AcrIF3 is a mimic of Cas8f HB and directly binds Cas2/3 to prevent it from being recruited (7,14).

AcrIF4 was identified in 2013 among the first five identified Acr proteins (15), and was found to bind the Csy complex in 2015 (16). While the structure of Csy-AcrIF4 complex has been solved in 2021 (17), the inhibition mechanism of AcrIF4 remains controversial. The structure of the Csy-AcrIF4 complex showed that AcrIF4 is wrapped between the spiral backbone formed by Cas7f and the tail composed of a Cas5f-Cas8f heterodimer of the Csy complex. Based on structural comparison, Gabel et al proposed that AcrIF4 will not prevent target DNA binding, which was further verified by the electrophoretic mobility shift assay (EMSA) in their study (17). However, a previous study in 2015 showed that AcrIF4 inhibits target dsDNA binding through *in vivo* experiments (16). Moreover, AcrIF4 binding has been proposed to prevent the rotation of Cas8f HB, which has not been experimentally verified (17). Taken together, the inhibition mechanism of AcrIF4 still remains enigmatic and also controversial about its effect on target DNA binding.

In this study, we first repeated the EMSA experiment of AcrIF4, which showed that the presence of AcrIF4 on the Csy complex does weakly inhibit target DNA binding. Notably, structural and biochemical studies revealed that AcrIF4 decreases target DNA binding through an approach very different from the canonical ones of competing with target DNA. The Csy-AcrIF4 complex can still bind target DNA, however, the Cas8f HB is held by AcrIF4 and does not rotate upon DNA binding to the Csy-AcrIF4 complex, which results in prevention of Cas2/3 recruitment and DNA cleavage. In all, our study presents an unprecedented mechanism in which AcrIF4 decreases target DNA binding to the Csy complex and inhibits Cas2/3 recruitment by anchoring the Cas8f HB.

## Results

### AcrIF4 decreases target DNA binding to the Csy complex

To clarify the present controversy about the effect of AcrIF4 on target DNA binding by the Csy complex, we repeated the EMSA experiment of AcrIF4 in the study of Gabel et al (17). The results showed that AcrIF4 in fact exhibits a weak inhibition on target DNA binding but does not completely prevent dsDNA binding even at a concentration 64 times as that of the Csy complex (Figure 1A). In comparison, both AcrIF1 and AcrIF13 can achieve a complete prevention at a concentration 4 times as that of the Csy (Figure 1A). This suggests that AcrIF4 is a only weak inhibitor of target DNA binding by the Csy complex, not as efficiently as AcrIF1 or AcrIF13 which directly hinders crRNA-DNA hybridization or acts as DNA mimic, respectively. To further explore the effect on dsDNA binding of the Csy complex by AcrIF4, we characterized the binding affinities of the Csy and Csy-AcrIF4 complex to dsDNA by EMSA assays. The results showed that the Csy-AcrIF4 complex exhibits a dsDNA binding *K*_d_ value about 2 times as that of the Csy complex (Figure 1B and Supplementary Figure S1), also suggesting a very weak inhibition by AcrIF4 binding. Interestingly, the binding of target DNA by the Csy-AcrIF4 complex was saturated only at a fraction DNA bound value of ∼0.75 (Figure 1B), suggesting an unstable binding between them. Taken together, the presence of AcrIF4 on the Csy complex results in a decreased binding to the target DNA, which is consistent with the previous results of *in vivo* experiment (16).

**Figure 1.**
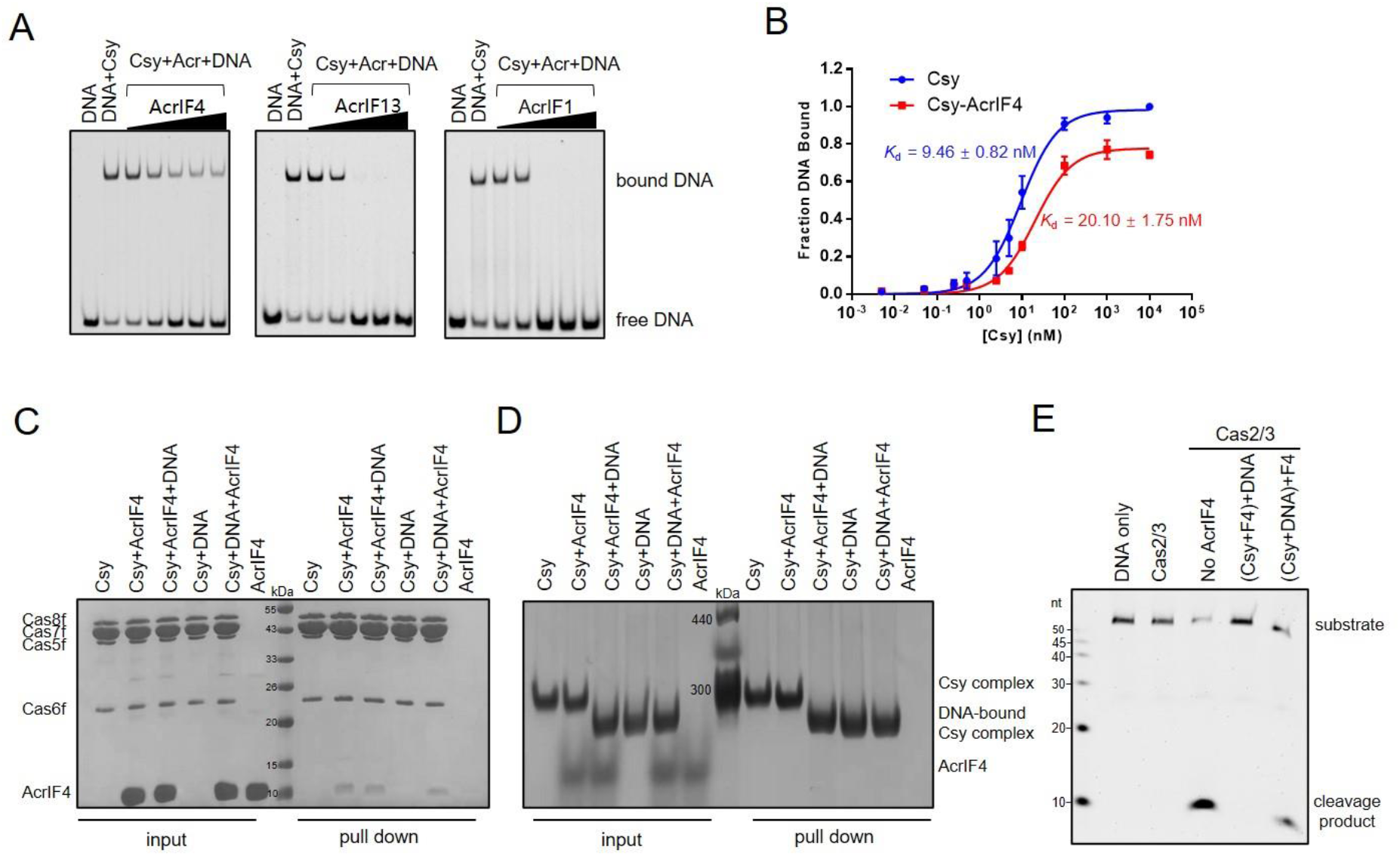
Binding of AcrIF4 to Csy decreases its DNA binding. (A) EMSA performed with dsDNA substrates shows that incubation with AcrIF4 results in reduced crRNA-guided DNA binding, but not as efficiently as with AcrIF1 and AcrIF13. Reactions were performed with 1.6 μM Csy, 0.1 μM dsDNA and Acr concentrations of 0.4, 1.6, 6.4, 25.6 and 102.4 μM following the order indicated by the black triangle. (B) The binding affinities of dsDNA to the apo Csy or the Csy-AcrIF4 complex tested by EMSA. Error bars represent SEM; n = 3. The *K*_D_ values are 9.46 ± 0.82 and 20.10 ± 1.75 nM for the Csy and Csy-AcrIF4, respectively. Raw data for these curves are shown in Figure S1. (C-D) Pull-down assay used to test the competition between AcrIF4 and dsDNA on the Csy complex. Reactions were performed with 6 μM Csy complex, 15 μM dsDNA and 180 μM AcrIF4 incubated in the required order (premix of Csy and DNA or premix of Csy and AcrIF4) for 30 min at 37°C, and then the mixtures were incubated with Ni-NTA beads for 30 min at 4°C. The reaction samples were run on SDS-PAGE (C) or native PAGE (D) after washing three times. (E) *In vitro* DNA cleavage reaction was performed with 0.4 μM Csy complex, 0.2 μM Cas2/3, and 0.04 μM 54-bp dsDNA (5’-FAM in the non-target DNA strand, NTS). AcrIF4 was added with concentrations of 6.4 μM following the order indicated in the figure.

### Csy-AcrIF4 complex binds target DNA but does not trigger its cleavage by Cas2/3

The above results suggested that while showing a decreased binding, Csy-AcrIF4 does bind target DNA as proposed by Gabel et al. (17). To confirm this, we performed a pull-down assay to investigate whether AcrIF4 and target DNA can co-exist on the Csy complex to form a ternary complex. After the pull-down assay, the samples were subjected to both SDS-PAGE gel and native PAGE, respectively, to detect the presence of dsDNA in the protein complex (Figure 1C, D). Due to the negative charge of dsDNA, the electrophoretic mobility of DNA-bound protein samples will run faster than the apo protein samples in the native PAGE, so that the migration position of the band can be used to determine whether there is bound DNA in the protein complex. The results showed that AcrIF4 and target DNA do not compete with each other on the Csy complex (Figure 1C, D). Target DNA can bind the preformed Csy-AcrIF4 complex, and AcrIF4 can also bind the preformed Csy-dsDNA complex. Up to now, most Acrs inactivate the CRISPR-Cas complex before target DNA binding, however, we recently found that AcrIF5 exhibits its inhibition capacity only after the Csy complex binds its target DNA (9). Therefore, we tested whether AcrIF4 inhibits the type I-F system before or after target DNA binding to the Csy complex. The results showed that preincubation of AcrIF4 and the Csy complex potently inhibits the activity of CRISPR-Cas system, but adding AcrIF4 after incubation of Csy and target DNA almost eliminates its inhibition capacity (Figure 1E). Taken together, AcrIF4 inhibits the CRISPR-Cas system by binding the Csy complex, and the Csy-AcrIF4 complex binds target DNA but does not trigger its cleavage.

### Overall structure of Csy-AcrIF4-dsDNA complex

To investigate why Csy-AcrIF4 binds DNA but does not trigger its cleavage by Cas2/3, we incubated Csy-AcrIF4 and target DNA, purified the complex to homogeneity (Supplementary Figure S2), and solved the structure of Csy-AcrIF4-dsDNA complex using single-particle cryoelectron microscopy (cryo-EM) at a resolution of 3.37 Å (Figure 2A and Supplementary Figure S3, and Supplementary Table S1). As shown in other Csy structures, the Csy complex is composed of an unequal stoichiometry of four different Cas proteins guided by a single 60-nt crRNA (Cas8f_1_:Cas5f_1_:Cas7f_6_:Cas6f_1_:crRNA_1_). Compared with the structure of Csy-AcrIF4 (PDB code: 7JZW), AcrIF4 binds the Csy complex at almost the same position in the Csy-AcrIF4-dsDNA complex, mainly surrounded by Cas8f and Cas5f subunits (Figure 2B). In the Csy-AcrIF4 structure, AcrIF4 also has contacts with Cas7.4f-7.6f subunits (Figure 2B). However, these contacts are all absent in the Csy-AcrIF4-dsDNA complex (Figure 2B), because of the elongation of the Csy backbone (to be introduced below). Consistently, densities for residues 49-52 of AcrIF4 are lacking in the Csy-AcrIF4-dsDNA structure, however, the same region is ordered and contacts Cas7.4f in the Csy-AcrIF4 structure (Figure 2C). Next, we compared the structures of Csy-AcrIF4-dsDNA and Csy-dsDNA (PDB code: 6NE0) complexes. Previous studies have revealed that, after target dsDNA binding, the Csy complex undergoes marked conformational changes compared to the apo Csy or that bound to a partially duplexed DNA target (7), which include clamping onto the dsDNA with the N-terminal segment of Cas8f, ∼18 Å elongation of the Csy backbone due to the movement of Cas8f-5f and hybridization between the target DNA and crRNA, and most importantly, a ∼180° rotation of the Cas8f HB (7). Compared to the structure of Csy-dsDNA, dsDNA binding to the Csy-AcrIF4 complex also induces movement of the N-terminal of Cas8f onto the dsDNA (Figure 2D, region 1), as well as elongation of the Csy backbone (Figure 2D, region 2 and Supplementary Figure S4). However, strikingly, the Cas8f HB is not rotated upon DNA binding to Csy-AcrIF4 (Figure 2D, region 3 and Figure 2E), and meanwhile no density for the displaced R-loop can be found in the Csy-AcrIF4-dsDNA structure (Figure 2F). The detailed differences between the structures of Csy-AcrIF4-dsDNA and Csy-dsDNA complexes will be discussed below.

**Figure 2.**
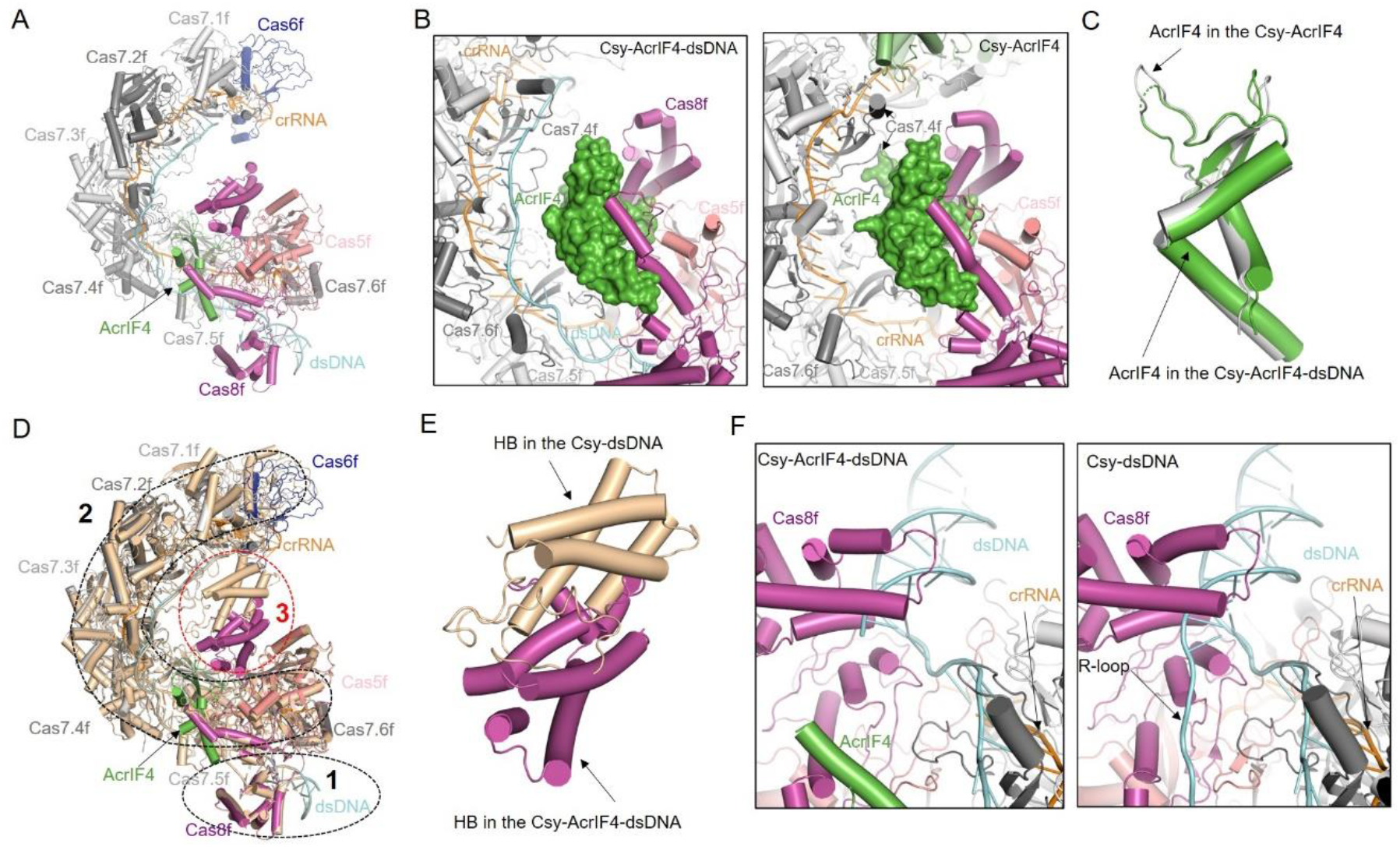
Overall cryo-EM structure of Csy-AcrIF4-dsDNA complex. (A) Atomic structure of Csy-AcrIF4-dsDNA in cartoon representation. (B) A close-up view of the AcrIF4 region in the Csy-AcrIF4-dsDNA (left) and Csy-AcrIF4 (right) complexes. (C)Comparison of the structures of AcrIF4 in Csy-AcrIF4-dsDNA and Csy-AcrIF4 complexes. (D) Comparison of the structures of Csy-AcrIF4-dsDNA (colored as in (A)) and Csy-dsDNA (colored in wheat). Region 1, the region of the N-terminal segment of Cas8f. Region 2, the region of the Csy backbone. Region 3, the region of Cas8f helical bundle (HB). (E)A close-up view of Region 3 in (D) showing the comparison of the structures of Cas8f HB in Csy-AcrIF4-dsDNA and Csy-dsDNA. (F) A close-up view of Region 1 in (D) showing the comparison of the R-loop regions in Csy-AcrIF4-dsDNA (left) and Csy-dsDNA (right) complexes.

### AcrIF4 binding prevents rotation of the Cas8f HB of Csy upon dsDNA binding

The most notable feature of Csy-AcrIF4-dsDNA structure is the lack of rotation of the Cas8f HB, compared to the structure of Csy-dsDNA (Figure 2E). Previous studies proposed that rotation of Cas8f HB of the Csy complex by dsDNA binding is dependent on R-loop formation (7), since this rotation was not observed in the structure of the Csy bound to a partially duplexed DNA which cannot form an R-loop (18). Interestingly, while a 54-bp fully duplexed DNA was used in our experiment, electron density only allows modelling of 39-nt in the target strand (TS) and 12-nt in the non-target strand (NTS) with no nucleotides in the R-loop region (Figure 2F). Close inspection of the structural alignment between Csy-AcrIF4-dsDNA and Csy-dsDNA revealed severe clash between AcrIF4 and the 9-nucleotide R-loop region of NTS (Figure 3A), suggesting a different path of the displaced NTS in the Csy-AcrIF4-dsDNA complex. Importantly, a previous study identified a positively charged channel formed by residues in Cas8f and Cas5f, named R-loop binding channel (RBC), which is important for the stabilization of the R-loop and therefore, dsDNA binding to the Csy complex (Figure 3B) (7). Importantly, AcrIF4 engages Cas8f R207/R219 and Cas5f R77 of the RBC (Figure 3C), and thus forces the R-loop to extend towards other positions. This may explain the lack of density of the R-loop, suggesting that it becomes flexible without the stabilization effect of the RBC. Moreover, rotation of the Cas8f HB will also expose several positively charged residues in Cas8f HB to form another part of the RBC (7), which is also inhibited in the Csy-AcrIF4-dsDNA complex. On the other hand, in addition to its role in Cas2/3 recruitment, rotation of the Cas8f HB upon DNA binding also contributes to a stable “locked” conformation of the crRNA-target-DNA duplex (7). In this conformation, the ‘thumbs’ of Cas7.2f and Cas7.3f fold over the target DNA strand and further bind the HB of Cas8f from one side, and their ‘webs’ interact with the Cas8f HB from the opposite site of the target DNA, thus completely locking the target DNA strand (Figure 3D). However, compared to the Csy-dsDNA structure, densities for the 5’-end five nucleotides of the TS are lacking in the Csy-AcrIF4-dsDNA structure (Figure 3D), suggesting that the absence of the locking effect of the Cas8f HB further destabilize the crRNA-DNA duplex in the 5’-end orientation of the TS of dsDNA. Taken together, we hypothesized that lack of both the stabilization of the R-loop by the RBC and the locking effect of Cas8f HB in the Csy-AcrIF4-dsDNA complex may increase the possibility of reannealing of the DNA duplex, thus decreasing the DNA binding ability of the Csy-AcrIF4 complex. To test this hypothesis, we repeated the EMSA experiment in Figure 1A with a dsDNA substrate containing a non-complementary ‘‘bubble”, which results in an R-loop incapable of reannealing (7,9). Consistent with our hypothesis, the Csy-AcrIF4 complex binds to the “bubble” dsDNA with a binding affinity similar as that of the Csy complex and can achieve a 100% binding (Figure 3E and Supplementary Figure S5). Moreover, distinct from AcrIF1 and AcrIF13, AcrIF4 does not decrease the binding of the “bubble” DNA by the Csy complex (Figure 3F). Taken together, our results indicated that the existence of AcrIF4 on the Csy complex blocks the RBC of the Csy complex and prevents the rotation of Cas8f HB upon dsDNA binding, thus reducing the R-loop stability and dsDNA binding ability of the Csy complex.

**Figure 3.**
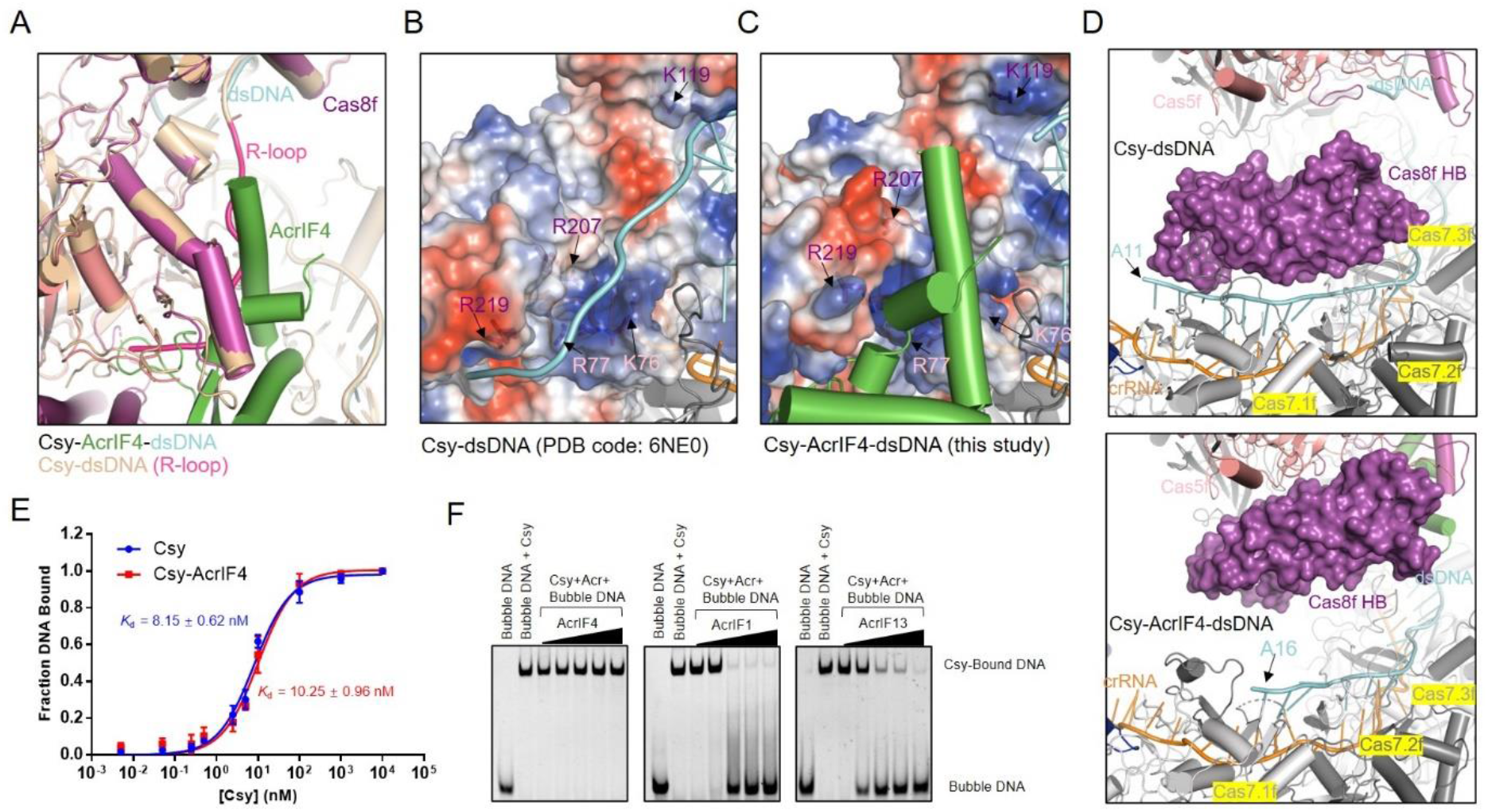
AcrIF4 binding to the Csy destabilize the R-loop formed by the NTS. (A)A close-up view of the R-loop region of the superimposition of the structures of the Csy-AcrIF4-dsDNA and Csy–dsDNA. The R-loop in Csy-dsDNA is colored in hot pink. (B-C) R-loop binding channel (RBC) in the structure of Csy-dsDNA (B) and Csy-AcrIF4-dsDNA (C). The Cas8f and Cas5f residues lining the RBC are shown as sticks and colored as the subunits in Figure 2A. The Cas8f and Cas5f subunits in two structures are shown in electrostatic model. (D) A close-up view of the 5’ end of TS in the Csy-dsDNA (top) and Csy-AcrIF4-dsDNA (bottom) complexes. The Cas8f HB in two structures are shown in surface model. (E) The binding affinities of “bubble” dsDNA by the apo Csy or the Csy-AcrIF4 complex tested by EMSA. Error bars represent SEM; n = 3. The *K*_D_ values are 8.15 ± 0.62 and 10.25 ± 0.96 nM for the Csy and Csy-AcrIF4, respectively. Raw data for these curves are shown in Figure S5. (F) EMSA assays showing that incubation with AcrIF4 has no effect on the binding of “bubble” DNA. Reactions were performed with 1.6 μM Csy, 0.1 μM “bubble” dsDNA and Acr concentrations of 0.4, 1.6, 6.4, 25.6 and 102.4 μM following the order indicated by the black triangle.

### AcrIF4 binding prevents Cas2/3 recruitment

It has been reported that rotation of the Cas8f HB of the Csy complex upon dsDNA binding will expose a “Cas2/3 recruitment helix” in Cas8f HB, which is essential for Cas2/3 recruitment and subsequent DNA cleavage (7). Therefore, we investigated whether the Csy-AcrIF4 complex has the ability to recruit Cas2/3 upon dsDNA binding through EMSA experiments. As shown in Figure 4A, the band of Csy-DNA complex shifted again with the addition of Cas2/3, indicating the formation of a ternary complex Csy-DNA-Cas2/3 (lanes 2–5). However, Cas2/3 recruitment was markedly inhibited when Csy-AcrIF4 was used instead of the Csy complex (Figure 4A, lanes 7-10). Notably, the Csy-AcrIF4 complex displays a reduced binding to dsDNA (compare lanes 2 and 7 in Figure 4A), which may also lead to less recruitment of Cas2/3. To avoid this, we also used the “bubble” dsDNA instead of fully duplexed dsDNA in the experiment shown in Figure 4A, which also showed that the presence of AcrIF4 prevents Cas2/3 recruitment by the Csy-dsDNA complex (Figure 4B). Since AcrIF4 can bind either the apo Csy complex or the Csy-dsDNA complex (Figure 1C, D), next, we tested different incubation orders with an AcrIF4 concentration gradient. The results showed that AcrIF4 displays potent inhibitory effect on Cas2/3 recruitment when Csy was preincubated with AcrIF4 (Figure 4C, lanes 4-6). However, AcrIF4 displayed no inhibitory effect on Cas2/3 recruitment when Csy and dsDNA were preincubated first (Figure 4C, lanes 7-9). This is consistent with the results of the *in vitro* DNA cleavage assay (Figure 1E), suggesting that prevention of Cas2/3 recruitment by binding the Csy is the major inhibition mechanism of AcrIF4.

**Figure 4.**
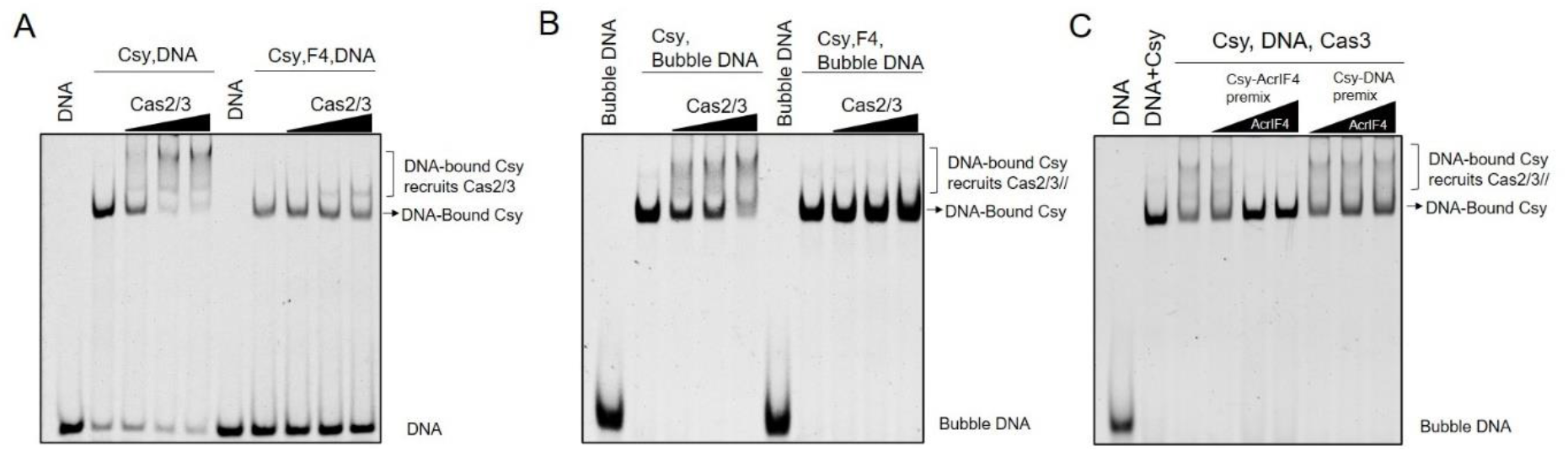
AcrIF4 prevents the recruitment of Cas2/3. (A-B) EMSA to test the Cas2/3 recruitment by the Csy-dsDNA (or Bubble dsDNA) complex or in the presence of AcrIF4. Reactions were performed with 1.6 μM Csy complex, 0.1 μM 54-bp dsDNA (5′-FAM in the target DNA strand), 25.6 μM AcrIF4, and concentrations of Cas2/3 were set as 0.2, 0.4 and 0.8 μM following the order indicated by the black triangle. Reactions were repeated independently three times with similar results. (C) The order of addition of AcrIF4 affects the inhibitory effect confirmed by an EMSA assay. Reaction was performed with 1.6 μM Csy complex, 0.8 μM Cas2/3 and 0.1 μM 54-bp dsDNA bubble (5′-FAM in the target DNA strand). AcrIF4 was added at the indicated order and with concentrations of 0.8, 3.2 and 12.8 μM. The experiments were repeated independently three times with similar results.

### Interaction with the Cas8f HB is essential for the inhibitory activity of AcrIF4

Notably, AcrIF4 has extensive interactions with the apo Csy complex with a ∼2580 Å^2^ buried surface, engaging the middle region and HB of Cas8f and Cas5f, as well as Cas7.4-7.6f (17). After DNA binding, the interfaces with Cas7f subunits are interrupted because of the elongation of the Csy backbone (Figure 2B). To investigate which interface is essential for the inhibitory effect of AcrIF4, we designed eight AcrIF4 mutants with mutations of residues involved in each interface (Figure 5A) and purified these proteins to homogeneity. Interestingly, pull-down assay showed that none of the mutations impair the interactions between AcrIF4 and the Csy complex (Supplementary Figure S6A and Figure 5B), suggesting that the Csy-AcrIF4 interface is too extensive to be broken by mutations in a single part of the interface. Consistently, all the AcrIF4 mutants can prevent Cas2/3 recruitment by the Csy-dsDNA complex as WT AcrIF4 in the EMSA experiments (Supplementary Figure S6B and Figure 5C). Because prevention of rotation of Cas8f HB is the inhibitory mechanism of AcrIF4, we wonder whether AcrIF4 will still retain its inhibitory capacity when its interface with the Cas8f HB is completely interrupted. However, AcrIF4 L39G/F54G, a mutant designed to interrupt its interaction with the Cas8f HB (Figure 5A, right panel), still inhibits recruitment of Cas2/3 (Figure 5C, left panel), suggesting that the Cas8f HB may still be held by the AcrIF4 mutant. To completely interrupt the interaction between AcrIF4 and the Cas8f HB, we further designed a Csy mutant with Cas8f R299G/R302A mutation in the Cas8f HB-AcrIF4 interface (Figure 5A, right panel). Pull-down assay showed that AcrIF4 L39G/F54G is still able to interact with this Csy mutant as WT Csy (Figure 5B). However, while the Csy mutant can recruit Cas2/3 upon dsDNA binding similarly as WT Csy, the AcrIF4 L39G/F54G mutant cannot inhibit either the Cas2/3 recruitment by the Csy mutant-dsDNA complex (Figure 5C, right panel) or DNA cleavage (Figure 5D). This suggested that although the AcrIF4 L39G/F54G mutant still binds the Csy mutant through the middle region of Cas8f and Cas5f, complete loss of the interaction with Cas8f HB renders AcrIF4 uncapable to hold the Cas8f HB in place and lose its inhibitory capacity. Moreover, it indicated that interaction between AcrIF4 and Cas8f HB is not essential for AcrIF4-Csy binding.

**Figure 5.**
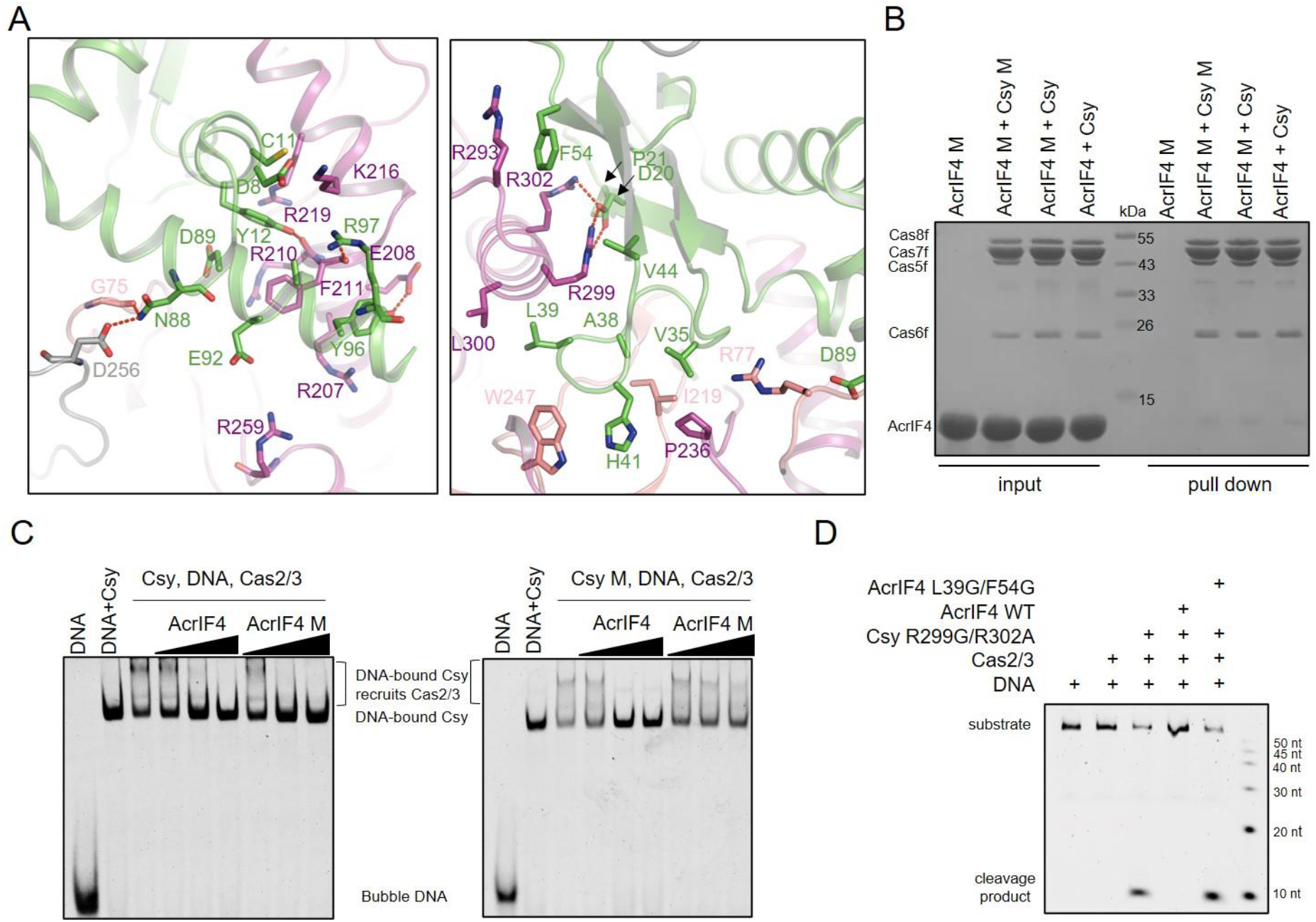
Interactions between AcrIF4 and the Cas8f HB are essential for the inhibition capacity of AcrIF4. (A) Detailed interactions between AcrIF4 and the Csy complex. The proteins are colored as in Figure 2A. (B) Mutations of the interface between the Cas8f HB and AcrIF4 do not affect the binding of AcrIF4 to the Csy complex. Reactions were performed with 6 μM Csy complex or its mutant, and 180 μM AcrIF4 or its mutant for 30 min at 37°C, and then the mixtures were incubated with Ni-NTA beads for 30 min at 4°C. Samples of input and pull-down were separated using SDS-PAGE after washing three times. Csy M, Cas8f R299G/R302A. AcrIF4 M, AcrIF4 L39/F54G. (C) EMSA to test the Cas2/3 recruitment by the Csy or its mutant (Cas8f R299G/R302A) in the presence of AcrIF4 or its mutant (AcrIF4 L39/F54G). Reactions were performed with 1.6 μM Csy complex, 0.1 μM 54-bp dsDNA bubble (5′-FAM in the target DNA strand), 0.8, 3.2, 12.8 μM μM AcrIF4, and concentrations of Cas2/3 were set as 0.8 μM following the order indicated by the black triangle. Reactions were repeated independently three times with similar results. Csy M, Cas8f R299G/R302A. AcrIF4 M, AcrIF4 L39/F54G. (D) *In vitro* DNA cleavage reaction was performed with 0.4 μM Csy mutant, 0.2 μM Cas2/3,6.4 μM AcrIF4 or its mutant and 0.04 μM 54-bp dsDNA (5′-FAM in the non-target DNA strand, NTS). The Csy complex and AcrIF4 were pre-incubated first, then dsDNA and Cas2/3 were added sequentially.

This also suggested that although the presence of AcrIF4 on the Csy complex keeps the R-loop away from the RBC on the middle region of Cas8f, the R-loop can still trigger the rotation of Cas8f HB when the AcrIF4-Cas8f HB interface is completely interrupted. Taken together, interaction with the Cas8f HB is essential for the inhibitory activity of AcrIF4 by holding the Cas8f HB in place.

## Discussion

Intense competition between bacteria and bacteriophages promotes microbial evolution, explaining why Acrs are highly different in their sequences, structures and inhibitory mechanisms (6). This also provides exciting resources for both researches into microbial life processes, and potential regulation tools of genome editing based on CRISPR-Cas systems. In this study, we describe the inhibitory mechanism of AcrIF4, to our knowledge the only known type I-F Acr that targets the apo Csy complex but does not use inhibition of target DNA binding as its major inhibitory mechanism. Our structural and functional data demonstrate that AcrIF4 prevents the Csy complex from completing the conformational changes necessary for Cas2/3 recruitment upon target dsDNA binding. Interestingly, AcrIF4 engages a very broad interface with the Csy complex to ensure that AcrIF4 can stably anchor the Cas8f HB. This resembles a “ship anchor” model, in which the extensive interface clamps AcrIF4 within the Csy complex, like an anchor wrapped in the sand, and the Cas8f HB is like a “ship” anchored by AcrIF4 and is prevented from moving (Figure 6).

**Figure 6.**
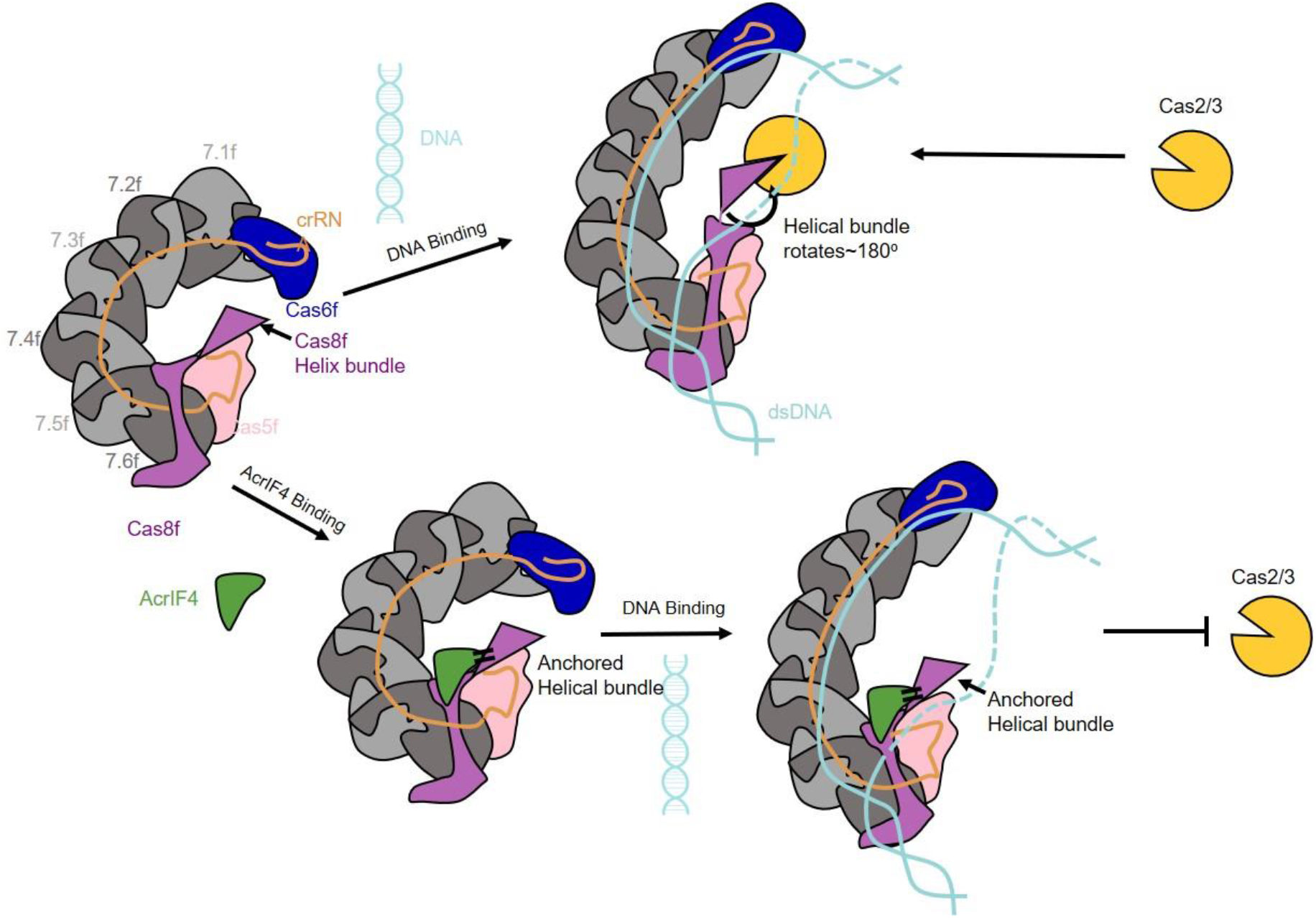
Working model of AcrIF4 in inhibition of the type I-F CRISPR-Cas system. AcrIF4 prohibits rotation of Cas8f HB caused by R-Loop formation. The extensive interfaces keep AcrIF4 firmly clamped in the Csy complex and the interaction between the Cas8f HB and AcrIF4 prevents Cas8f HB from moving.

Previous studies are controversial about whether the binding of AcrIF4 affects DNA binding of the Csy complex. While *in vivo* assays showed the presence of AcrIF4 decreases DNA binding to the Csy (16), the EMSA result by Gabel et al. showed no effect of AcrIF4 on this binding (17). Our EMSA results indicated that AcrIF4 does weakly inhibit DNA binding to the Csy complex. However, the binding site of AcrIF4 on the Csy complex neither conflicts with the hybridization site of the target DNA strand on Cas7f subunits nor shields the PAM recognition region on Cas8f. This is reflected by the fact that Csy-AcrIF4 can still bind dsDNA to form a ternary complex, which is significantly different from the canonical competition approaches utilized by AcrIF1/2/6/7/8/9/10/13/14/24 (11-13,17-23). We found that AcrIF4 shields some residues of the RBC and also disrupts the Cas8f HB part of RBC by preventing rotation of the Cas8f HB. The lack of both the stabilization of the R-loop by the RBC and the locking effect on target DNA strand by the rotated Cas8f HB may collectively decrease the DNA binding to the Csy-AcrIF4 complex.

Up to now, out of the 15 Acrs including AcrIF4 with known inhibition mechanisms, only AcrIF4 and AcrIF5 can coexist with target DNA, which is bound at the right place, on the Csy complex. AcrIF4 is also an Acr that utilizes the conformational change of the Cas8f HB to exert its inhibitory effect. Unlike AcrIF5 which specifically binds the Csy-dsDNA complex to compete off the Cas8f HB and make it flexible, AcrIF4 sticks the Cas8f HB tightly to prevent its rotation upon dsDNA binding. In contrary to AcrIF5 which functions only after the formation of Csy-dsDNA complex, AcrIF4 functions by binding the Csy complex before target DNA binding. Therefore, elucidation of the mechanism of AcrIF4 also has evolutionary implications that targeting the same procedure of host immunity can be executed in different ways by phage proteins. In all, our study reveals an unprecedented anti-CRISPR mechanism and highlights the functional diversity of Acr proteins.

### Experimental Procedures

#### Protein expression and purification

The full-length AcrIF4 gene was synthesized by GenScript and amplified by PCR and cloned into a modified pET28a vector which encodes a SUMO protein to produce a His-SUMO tagged fusion protein with a Ulp1 peptidase cleavage site between SUMO and the target protein. The mutants of AcrIF4 or the Csy complex were generated by two-step PCR and were subcloned, overexpressed, and purified in the same way as the wild-type protein. The AcrIF4 protein was expressed in *E. coli* strain BL21 (DE3) and induced by 0.2 mM isopropyl-β-D-thiogalactopyranoside (IPTG) when the cell density reached an OD_600nm_ of 0.8. After growth at 16 °C for 12 h, the cells were harvested, re-suspended in lysis buffer (300mM NaCl, 50 mM Tris-HCl pH 8.0, 30 mM imidazole and 1 mM PMSF) and lysed by sonication. The cell lysate was centrifuged at 20,000 g for 45 min at 4 °C to remove cell debris. The supernatant was applied onto a self-packaged Ni-NTA affinity column (2 mL High Affinity Ni-NTA Resin; GenScript) and contaminant proteins were removed with wash buffer (300mM NaCl, 50 mM Tris-HCl pH 8.0, 30 mM imidazole). The AcrIF4 protein was eluted from the Ni column after reaction with Ulp1 peptidase at 18°C for 2 h. The eluant was concentrated and further purified using a Superdex-75 (GE Healthcare) column equilibrated with a buffer containing 10 mM Tris–HCl pH 8.0, 200 mM NaCl, and 5 mM DTT. The purified protein was analyzed by SDS– PAGE. The Csy complex and Cas2/3 were cloned, overexpressed and purified as described previously (13). To obtain the Csy-AcrIF4 complex, the AcrIF4 gene was cloned into the pACYCDuet-1 vector together with the gene that encodes the pre-crRNA, and the Csy-AcrIF4 complex can be purified using the same purification method as the Csy complex.

The Csy-AcrIF4-dsDNA complex for cryo-EM study was prepared as follows. The Csy-AcrIF4 complex was incubated with dsDNA in a molecular ratio of 1: 2.5 (Csy-AcrIF4: dsDNA). Then the Csy-AcrIF4-dsDNA complex was separated using a Superdex-200 (GE Healthcare) column equilibrated with a buffer containing 10 mM Tris–HCl pH 8.0, 200 mM NaCl, and 5 mM DTT. Purified proteins were finally flash-frozen in liquid nitrogen.

#### Cryo-EM sample preparation and data acquisition

Aliquots (4 μL) of Csy-AcrIF4-dsDNA complex (2.5 mg/mL) were applied to carbon grids (Quantifoil 300-mesh Au R1.2/1.3, Micro Tools GmbH, Germany). The grids were blotted for 1.5 s and plunged into liquid ethane in 100% humidity at 8 °C with Mark IV Vitrobot (Thermo Fisher Scientific). 839 raw movie stacks were collected using a Titan Krios microscope (Thermo Fisher Scientific) operated at 300 kV by a K3 Summit direct electron detector using SerialEM software at a nominal magnification of 29,000 × in super-resolution mode with a total dose of 50 e/Å^2^ and exposure time of 3 s and the defocus range were set from −1.3 μm to −1.8 μm.

#### Image processing

In general, movies were motion-corrected using MotionCor2 (24). Gctf (25) was used to determine the contrast transfer function (CTF) parameter and produce the CTF power spectrum on basis of summed micrographs. Particles were auto-picked on dose-weighted micrographs using Laplacian of Gaussian in RELION 3.1 (26). Briefly, about 2,000 particles were selected and generated 2D averages templates for particle auto-picking and 621,597 particles were extacted from manually selected micrographs. The previously reported model of Csy-AcrIF14-dsDNA complex (PDB: 7ECW) was low-pass filtered to 40 Å for initial model. All extracted particles using a box size of 240 and a binned pixel size of 7.76 Å, were subjected to two rounds of 2D classification and 3D classification using C1 symmetry and 238,361 particles were retained. 117,510 selected particles were subjected to 3D auto-refinement without symmetry and performed an overall resolution of 3.37 Å. Meanwhile, Cas8f and Cas6f were signal subtracted based on the refined map with soft mask, then particles were subjected to 3D auto-refine and 3D classification without image alignment. Finally, 49,498 particles and 93,188 particles were separately performed to 3D auto-refine and yielded a map at 3.59 Å and 3.96 Å resolution. The composite map of Csy-AcrIF4-dsDNA complex was generated in UCSF Chimera (27) using ‘vop maximum’ command and used for model building and refinement. All reported resolutions are based on the gold-standard FSC=0.143 criteria, and the final FSC curves were corrected for the effect of a soft mask by using high-resolution noise substitution. The final density maps were sharpened by B-factors calculated with the RELION post-processing program. The final maps for model building and figure presentation were performed using DeepEMhancer (28). Local resolution map was calculated using ResMap (29). Further information for all samples is provided in Supplementary Table S1.

#### Model Building, Refinement and Validation

Atomic models of the Csy-AcrIF4-dsDNA complex were modelled using the PDB 6NE0. AcrIF4 was modelled using the PDB 7JZW. The initial model was manually refined using Coot (30) and subjected to real_space_refinement using PHENIX (31). All the figures were created in the PyMOL software.

#### Double-stranded DNA preparation

For EMSA and *in vitro* DNA cleavage assay, various 5′-end FAM labeled single-stranded DNA molecules were synthesized from Sangon, Shanghai, and were hybridized with their complementary unlabeled single-stranded DNA with a molar ratio of 1:1.5 to obtain double-stranded DNA. Specifically, dsDNA molecules used in pull-down assay were generated through the same method described above except that both strands were unlabeled and added with a 1: 1 ratio.

Target DNA strand (54 bp; 5’-FAM fluorescein labeled or unlabeled) GGAAGCCATCCAGGTAGACGCGGACATCAAGCCCGCCGTGAAGGTGCAGCTGCT

Non-Target DNA strand (54 bp; 5’-FAM fluorescein labeled or unlabeled) AGCAGCTGCACCTTCACGGCGGGCTTGATGTCCGCGTCTACCTGGATGGCTTCC

Non-Target DNA strand of the dsDNA bubble (54 bp; unlabeled) AGCAGCTGCACCAAGTGCCGCCGCTTGATGTCCGCGTCTACCTGGATGGCTTCC

#### Electrophoretic mobility shift assay

##### Binding of Acr to the Csy complex affects dsDNA binding

Duplexed DNA was prepared as above with 5’ FAM label at the target strand. Reactions were performed by incubating 1.6 μM Csy complex with 0.4, 1.6, 6.4, 25.6 and 102.4 μM Acr for 30 min at 37°C in a buffer containing 20 mM HEPES pH 7.5, 100 mM KCl, 5% glycerol, and 1 mM TCEP. And then 0.1 μM dsDNA (or dsDNA bubble) was added and incubated for another 30 min at 37°C. The mixtures were separated using 5% native polyacrylamide gels and visualized by fluorescence imaging.

##### Cas2/3 recruitment

100 nM dsDNA or 0.8, 3.2, 12.8 μM AcrIF4 was pre-incubated with 1.6 μM Csy complex at 37 °C in the reaction buffer (20 mM HEPES pH 7.5, 100 mM KCl, 5% glycerol, 1 mM TCEP) for 30 min, and then AcrIF4 or dsDNA was added and incubated for another 30 min. Afterwards, Cas2/3 was added to a final concentration of 0.8 μM. The reaction was further incubated for 10 min. Products of the reaction were separated using 5% native polyacrylamide gels and visualized by fluorescence imaging.

##### Binding between dsDNA and the Acr-bound Csy complex

The apo or the AcrIF4-bound Csy complex was incubated in a concentration gradient (0, 0.005, 0.05, 0.25, 0.5, 2.5, 5, 10, 100, 1000, 10000 nM) with 16 nM 54 bp-dsDNA or bubble dsDNA (5’-FAM in the target strand). Binding reactions were conducted at 37°C for 30 min in the buffer containing 20 mM HEPES pH 7.5, 100 mM KCl, 5% glycerol, and 1 mM TCEP. Products of the reaction were separated using 5% native polyacrylamide gels and visualized by fluorescence imaging. The fluorescence signal was measured using ImageJ(32).

#### Ni-column pull-down assay

6 μM Csy complex, 15 μM dsDNA and 180 μM AcrIF4 or mutants were incubated in the required order (Csy and DNA first or Csy and AcrIF4 first) for 30 min at 37°C, and then the mixtures were incubated with Ni-NTA beads for 30 min at 4°C. The buffer containing 300 mM NaCl, 50 mM Tris-HCl pH 8.0, 30 mM imidazole was used to wash the beads. Samples of input and pull-down were separated using SDS-PAGE and native PAGE after washing three times.

##### *In vitro* DNA cleavage assay

First, 6.4 μM AcrIF4 or mutant and 0.4 μM Csy complex were incubated at 37°C for 30 min in the reaction buffer containing 20 mM HEPES pH 7.5, 100 mM KCl, 5% glycerol, and 1 mM TCEP. And then 0.04 μM dsDNA (5’-FAM in the not-target strand) was added and incubated for another 30 min. The incubation order of DNA and AcrIF4 is adjusted according to the purpose of the experiment. Afterwards, Cas2/3 was added to a final concentration of 0.2 μM, along which 5 mM MgCl_2_, 75 μM NiSO_4_, 5 mM CaCl_2_, and 2 mM ATP were added into the buffer. The reaction was further incubated for 30 min and quenched with 1% SDS and 50 mM EDTA. The products were separated by electrophoresis over 14% polyacrylamide gels containing 8 M urea and visualized by fluorescence imaging.

## Data Availability

Cryo-EM reconstruction of Csy-AcrIF4-dsDNA has been deposited in the Electron Microscopy Data Bank under the accession numbers EMD-33837. The coordinate for atomic model of Csy-AcrIF4-dsDNA has been deposited in the Protein Data Bank under the accession number 7YHS.

## Acknowledgements

We thank J. Lei and F. Yang at Tsinghua University for data collection. We acknowledge support with cryo-EM facilities from the Tsinghua University Branch of the China National Center for Protein Sciences (Beijing). This work was supported by the National key research and development program of China (2019YFC1200500 and 2019YFC1200502), the National Natural Science Foundation of China (32000901 and 32171274), Beijing Nova program and the Fundamental Research Funds for the Central Universities (XK1802-8).

## Conflict of interest statement

The authors declare no competing interests.

## Author Contributions

**Zhengyu Gao**: Methodology, Investigation. **Zihao Ge**: Methodology, Investigation. **Laixing Zhang**: Software, Data analysis. **Hao Wang**: Investigation. **Yourun Yue**: Investigation. **Zhuobing Jiang**: Investigation. **Xin Wang**: Investigation. **Chenying Xu**: Investigation. **Yi Zhang**: Supervision. **Maojun Yang**: Supervision. **Yue Feng**: Conceptualization, Methodology, Writing, Review & Editing.

## Notes

### Competing Interest Statement

The authors have declared no competing interest.

