## Supplementary Table S1 and Figure S1-S6 for "Structural and mechanistic insights into inhibition of the type I-F CRISPR-Cas surveillance complex by AcrIF4"

**Table S1. Cryo-EM data collection, refinement, and validation statistics**

| | Csy-AcrIF4-dsDNA-overall | Csy-AcrIF4-dsDNA-Cas8f | Csy-AcrIF4-dsDNA-Cas6f |
|---|---|---|---|
| **Data collection and processing** | | | |
| EMDB code | EMD-33837 | | |
| PDB code | 7YHS | | |
| Electron microscopy | | Titan Krios | |
| Camera | | K3 | |
| Magnification | | 29000× | |
| Voltage | | 300 kV | |
| Defocus range (μm) | | -1.3 to -1.8 | |
| Electron exposure (e-/Å$^2$) | | 50 | |
| Pixel size (Å) | | 0.97 | |
| Exposure rate (e$^-$/A$^2$/sec) | | 20 | |
| Number of frames per movie | | 32 | |
| Automation software | | SerialEM | |
| Micrographs collected | | 839 | |
| Data processing software | | RELION 3.1 | |
| Symmetry imposed | *C*1 | *C*1 | *C*1 |
| Total extracted particles | | 621,597 | |
| Total refined particles | 117,510 | 49,498 | 93,188 |
| Final particles | 117,510 | 49,498 | 93,188 |
| Map resolution (FSC=0.143/Å) | 3.37 | 3.59 | 3.96 |
| Local resolution range (Å) | 2.4-4.4 | 2.4-4.4 | 3.2-5.4 |
| **Refinement** | | | |
| Initial Model (PDB code) | 6NE0 | | |
| Refinement Package | Phenix 1.19 (real space refinement) | | |
| Model resolution (FSC=0.5/Å) | 3.43 | | |
| Map sharpening B factor (Å$^2$) | -20 | | |
| Map CC | 0.67 | | |
| Model composition | | | |
| Non-hydrogen atoms | 24,594 | | |
| Protein residues | 2949 | | |
| Nucleotides | 109 | | |
| *B* factors (Å$^2$) | | | |
| Protein | 28.17 | | |
| Nucleotides | 53.28 | | |
| R.m.s. deviations | | | |
| Bond lengths (Å) | 0.005 | | |
| Bond angles (°) | 0.876 | | |
| **Validation** | | | |
| MolProbity score | 2.42 | | |
| Clashscore | 5.43 | | |
| Poor rotamers (%) | 6.94 | | |
| C-beta deviations | 0 | | |
| CaBLAM outliers | 5.80 | | |
| Ramachandran plot (%) | | | |
| Favored | 91.87 | | |
| Allowed | 8.06 | | |
| Outlier | 0.07 | | |

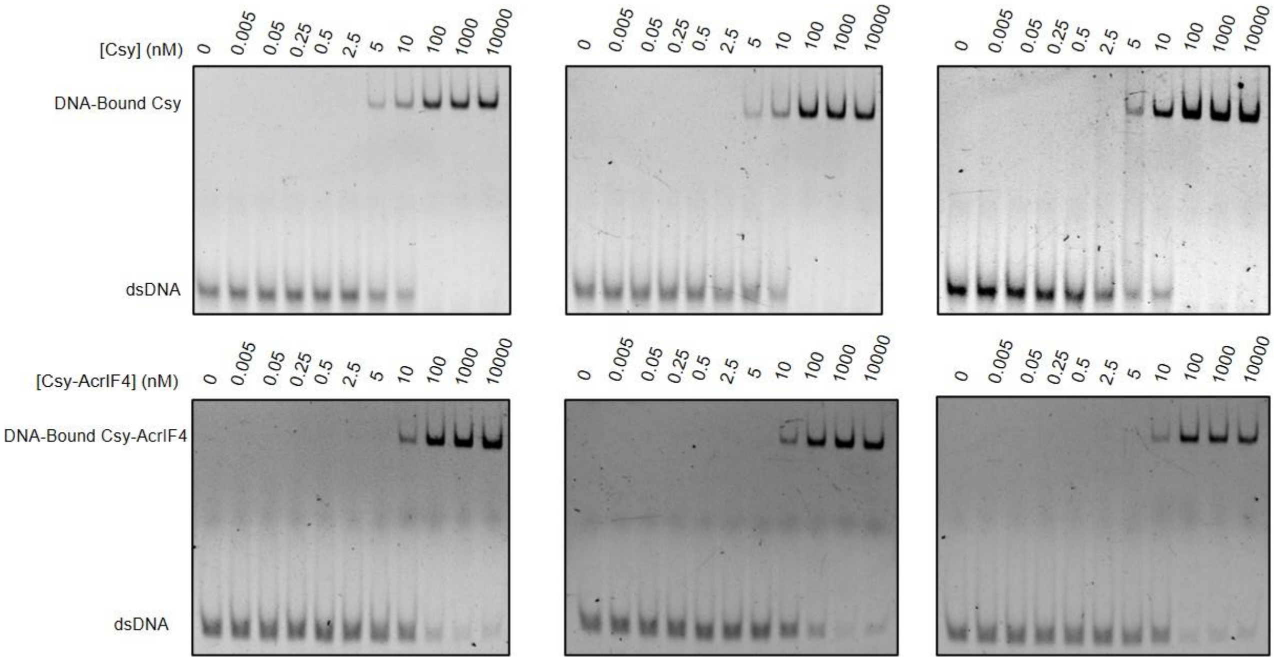


**Figure S1 Binding between dsDNA and the AcrIF4-bound or apo Csy complex**

DNA binding assays were performed by incubating a concentration gradient (0, 0.005, 0.05, 0.25, 0.5, 2.5, 5, 10, 100, 1000, 10000 nM) of the Csy (or Csy-AcrIF4) complex with 16 nM of 54 bp dsDNA (5′-FAM in the TS).

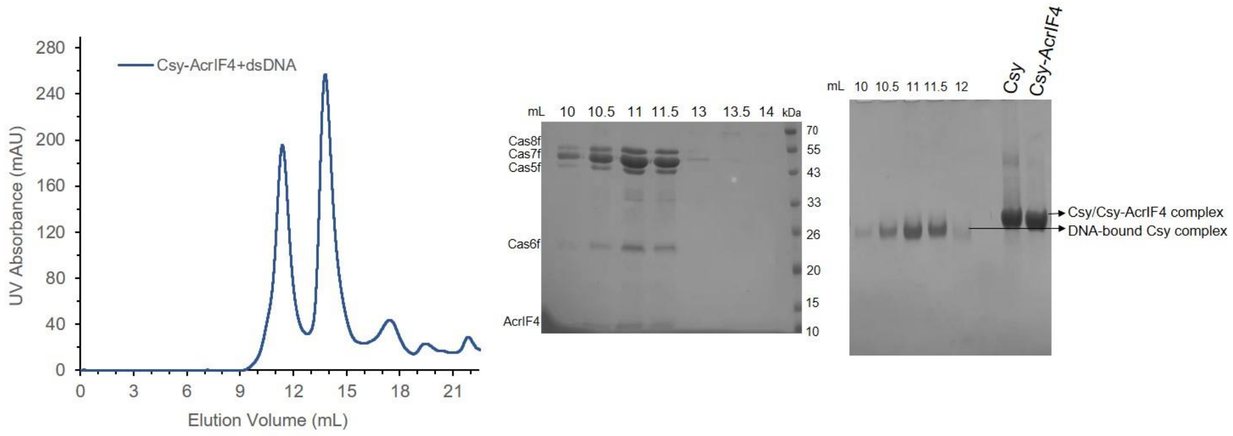


**Figure S2 dsDNA co-elutes with the Csy-AcrIF4 complex**

Gel filtration profiles of the mix of Csy-AcrIF4/dsDNA. The UV absorbance at 280 nm is shown. The SDS-PAGE and native PAGE of the peak fractions are shown on the right.

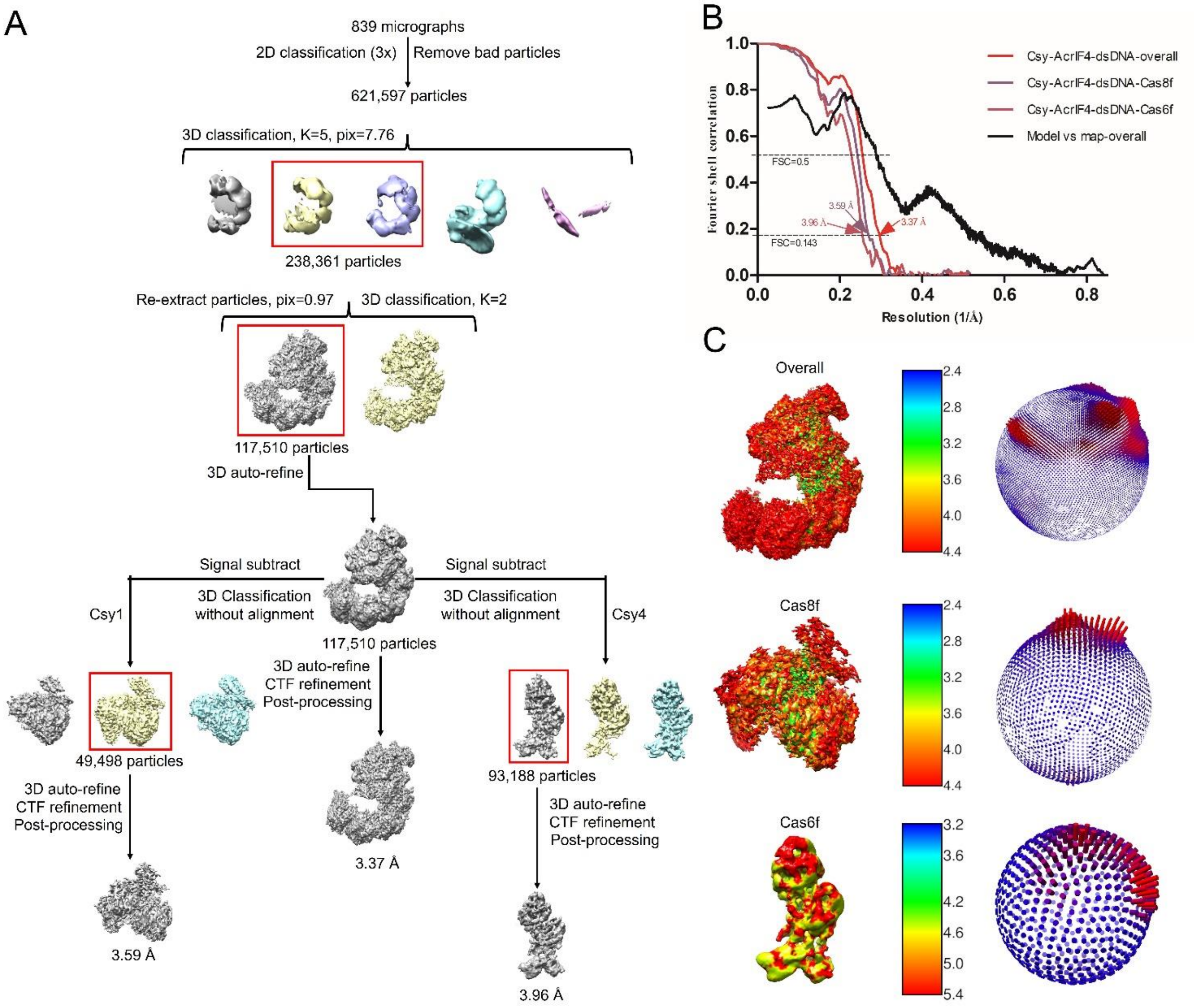


**Figure S3 Cryo-EM image processing for the Csy-AcrIF4-dsDNA complex**

(A) Representative data processing of Csy-AcrIF4-dsDNA complex. 3D classification, signal subtraction, 3D-auto refinement, CTF Refinement and post-processing were performed with subregion of Cas8f and Cas6f in RELION 3.1.

(B) Representative the gold-standard Fourier Shell Correlation (FSC=0.143) curves and model vs map curve (FSC=0.5) of Csy-AcrIF4-dsDNA complex, Cas8f and Cas6f.

(C) Local resolution estimation and particle orientation distributions of the Csy-AcrIF4-dsDNA complex, Cas8f and Cas6f.

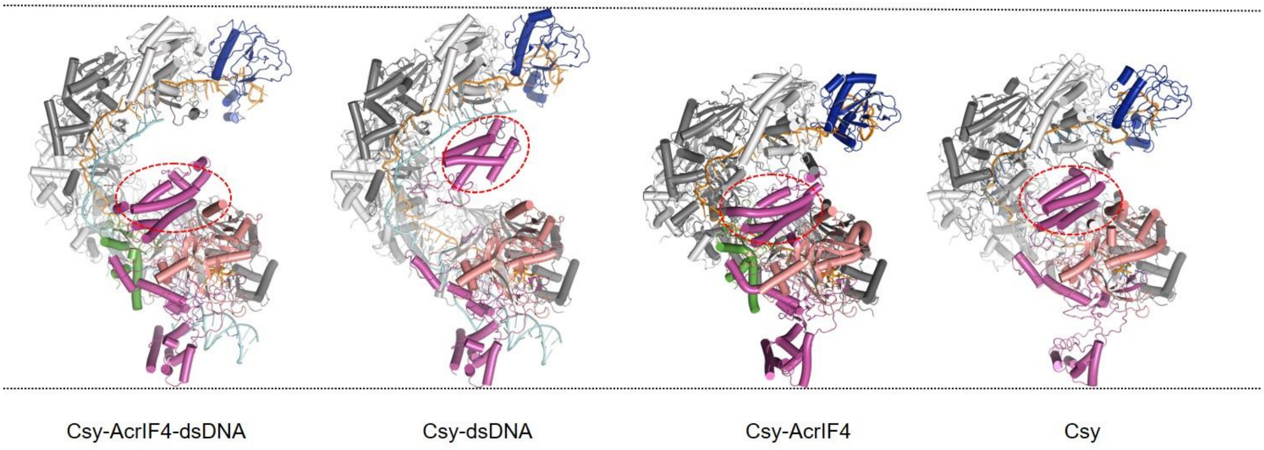


**Figure S4 Side-by-side comparison of the Csy-AcrIF4-dsDNA, Csy-dsDNA, Csy-AcrIF4 and Csy structures**

Side-by-side comparison of the Csy-AcrIF4-dsDNA (this study), Csy-dsDNA (PDB code: 6NE0), Csy-AcrIF4 (PDB code: 7JZW) and Csy (PDB code: 6B45) structures. The Cas8f HB is marked in a circle.

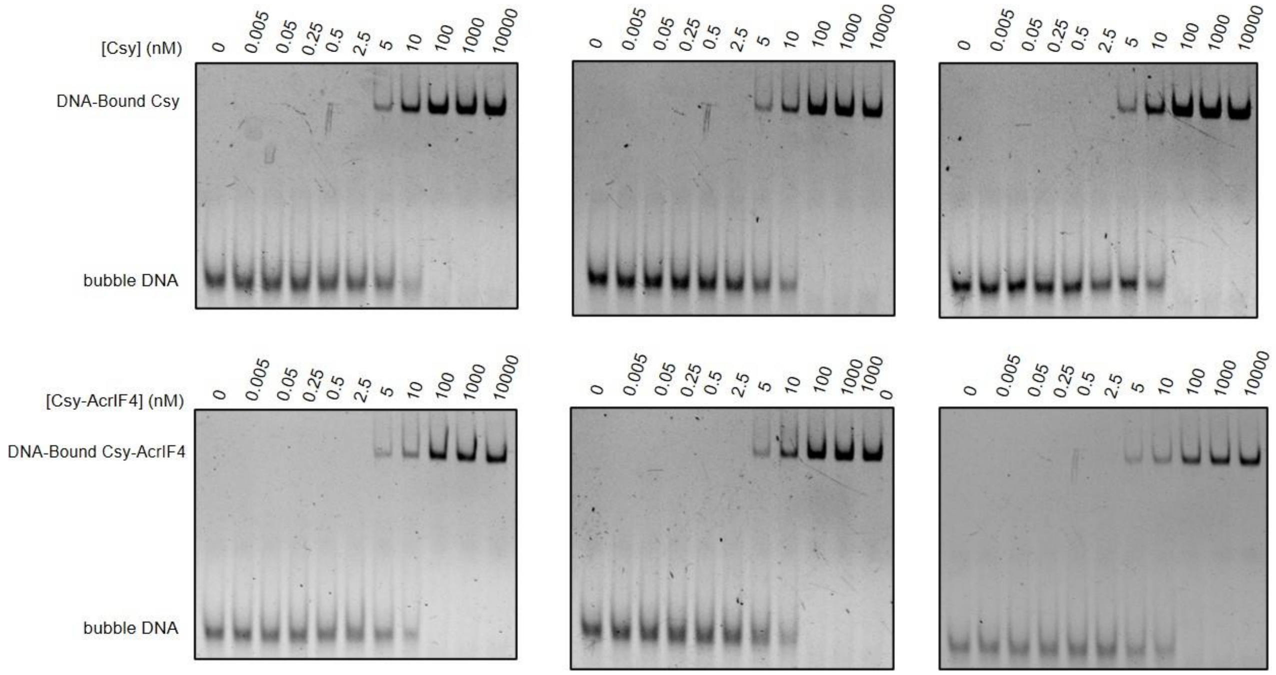


**Figure S5 Binding between bubble dsDNA and the AcrIF4-bound or apo Csy complex**

DNA binding assays were performed by incubating a concentration gradient (0, 0.005, 0.05, 0.25, 0.5, 2.5, 5, 10, 100, 1000, 10000 nM) of the Csy (or Csy-AcrIF4) complex with 16 nM of 54 bp dsDNA bubble (5′-FAM in the TS).

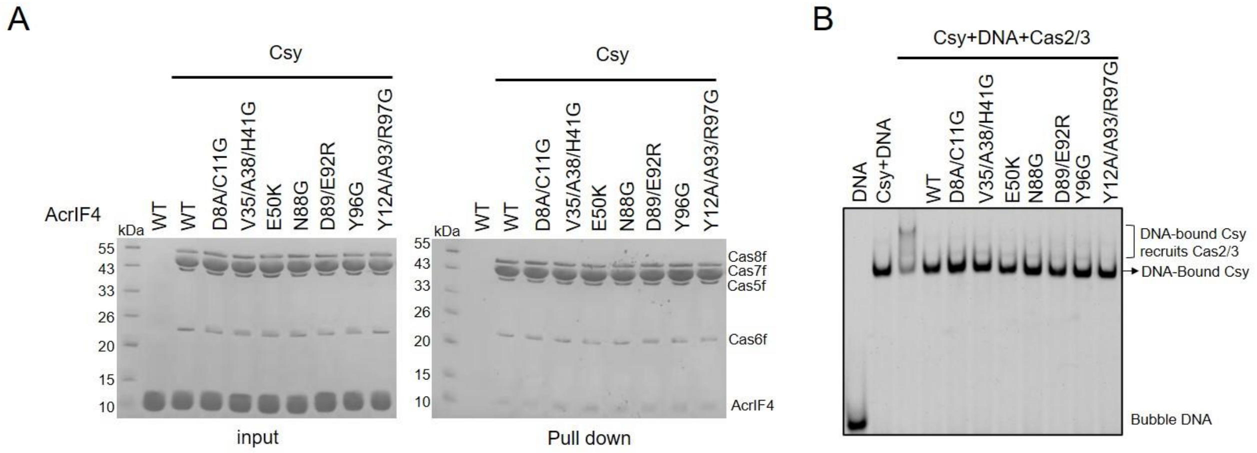


**Figure S6 AcrIF4 mutations on a specific interface do not result in marked Csy binding defect**

(A) Reactions were performed with 6 μM Csy complex and 180 μM AcrIF4 or its mutants for 30 min at 37℃, and then the mixtures were incubated with Ni-NTA beads for 30 min at 4℃. Samples of input and pull-down were separated using SDS-PAGE after washing three times.

(B) Mutations of the interface residues of AcrIF4 did not affect its inhibition capacity of Cas2/3 recruitment. Reactions were performed with 1.6 μM Csy complex, 0.1 μM 54-bp bubble dsDNA (5′-FAM in the target DNA strand), 3.2 μM AcrIF4, and 0.8 μM Cas2/3. Reactions were independently repeated three times with similar results.
